# CO_2_ protects cells from iron-Fenton oxidative DNA damage in E. coli and humans

**DOI:** 10.1101/2024.08.26.609766

**Authors:** Aaron M. Fleming, Justin C. Dingman, Cynthia J. Burrows

## Abstract

Whereas hydroxyl radical is commonly named as the Fenton product responsible for DNA and RNA damage in cells, here we demonstrate that the cellular reaction generates carbonate radical anion due to physiological levels of bicarbonate. Analysis of the metabolome, transcriptome and, in human cells, the nuclear genome shows a consistent buffering of H_2_O_2_-induced oxidative stress leading to one common pathway, namely guanine oxidation. Particularly revealing are nanopore-based studies of direct RNA sequencing of cytosolic and mitochondrial ribosomal RNA along with glycosylase-dependent qPCR studies of oxidative DNA damage in telomeres. The focusing of oxidative modification on one pathway is consistent with the highly evolved base excision repair suite of enzymes and their involvement in gene regulation in response to oxidative stress.

## Introduction

The necessity of iron for life and the discovery of H_2_O_2_ as a signaling molecule raises the question of how cells manage Fenton chemistry (*1*). It is widely believed that the reaction of Fe(II) + H_2_O_2_ generates powerful oxidants such as hydroxyl radical (HO^•^) or an Fe(IV)-oxo (ferryl, FeO^2+^) species, depending on pH and ligands (*2*). Both HO^•^ and FeO^2+^ react at near diffusion-controlled rates with all biomolecules, and though concentrations of both the labile iron pool (∼5 μM) and cellular H_2_O_2_ (0.1-0.5 μM) are very low (*3, 4*), any DNA damage resulting from HO^•^ oxidation would place an enormous burden on the DNA repair system that is tasked with locating and correcting the various forms of base and ribose damage and resulting repair intermediates (Scheme 1). Ultimately, unrepaired DNA lesions contribute to muta
genicity which is a key feature of cancer, neurological and age-related disorders (*5*). The burden of DNA repair could be lessened if metabolic H_2_O_2_ generated a weaker, more selective, reactive oxygen species (ROS) during reaction with cellular iron. Here we demonstrate that CO_2_ and its hydrated form bicarbonate, HCO_3_^-^, not only buffer the pH of cells but also transform the Fenton reaction into one that targets only the guanine (G) base in nucleic acids.

**scheme 1.**
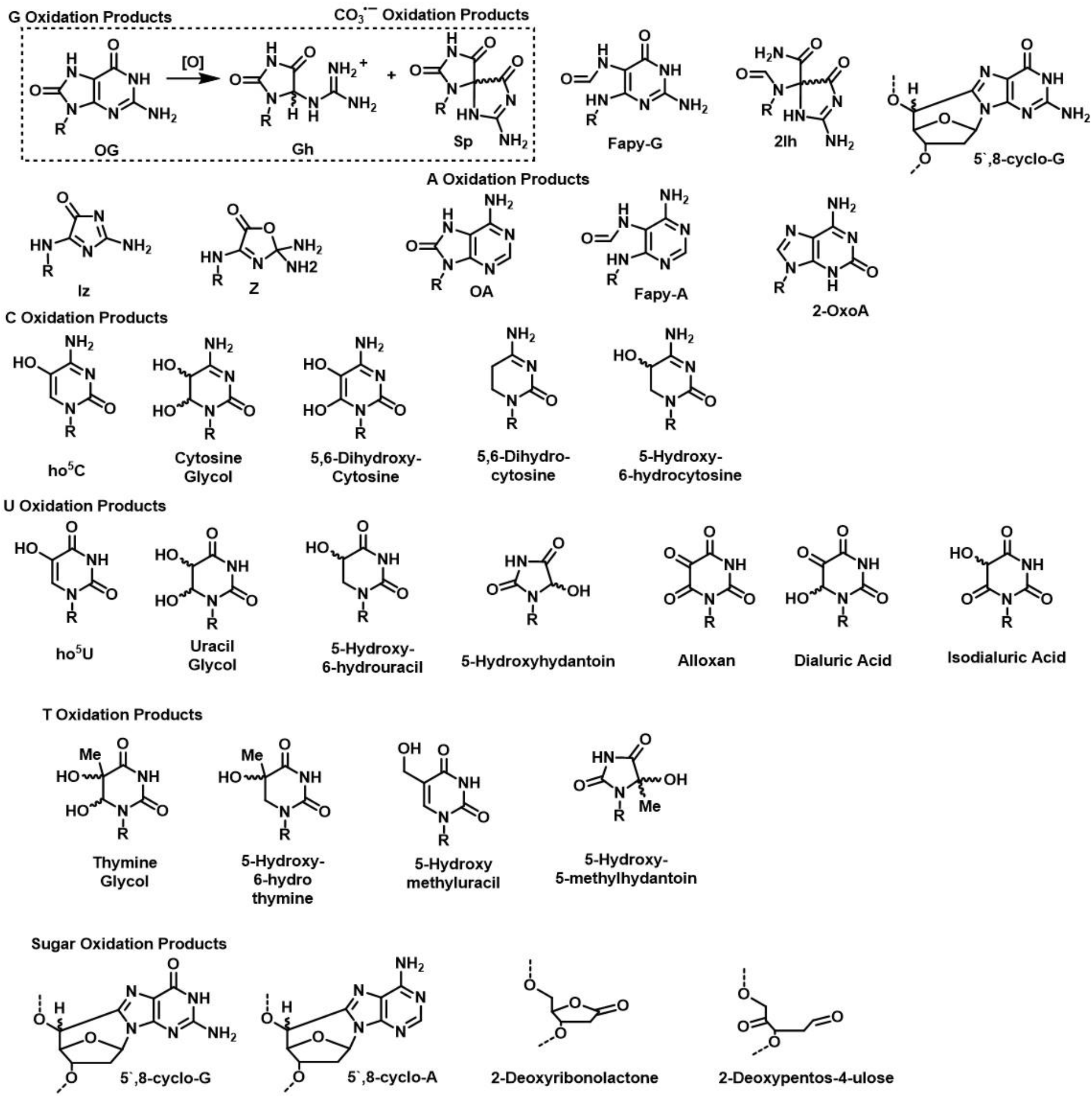
products of ho^•^ or co_3_^•-^ (dashed box) oxidative modification of nucleic acids.

In 2019, Meyerstein and coworkers demonstrated that aqueous Fenton reactions at pH 7 in the presence of bicarbonate result in an inner-sphere reaction of H_2_O_2_ with a Fe(II)-carbonate complex releasing carbonate radical anion (CO_3_^•-^; Eqn.1), not hydroxyl radical (*6, 7*). Accordingly, they proposed that the oxidant in biological Fenton reactions should be CO_3_^•-^, as long as sufficient HCO_3_^-^ is present.

Our initial model study also demonstrated that in the presence of 25 mM NaHCO_3_ which is a typical concentration for bicarbonate in eukaryotic cells, the in vitro Fenton oxidation of guanosine led almost entirely to the formation of 8-oxo-7,8-dihydro-2’-deoxyguanosine (OG) (*8*). Conversely, the classical Fenton reaction in phosphate buffer generates at least four other products arising from H^•^ abstraction by HO^•^. Importantly, carbonate radical anion is a sufficiently powerful oxidant to remove one electron from purines (A or G), but not pyrimidines, leading to purine radical cation formation in nucleic acids (*9*), and because of efficient charge transport in DNA (*10*), adenosyl radical cations are predicted to undergo hole transfer to form G^•+^, leading to OG as the major outcome (*11, 12*). Thus, the presence of bicarbonate in cellular buffers should focus the Fenton reaction on G by creating OG lesions, which are only mildly mutagenic and readily recognized by base excision repair glycosylases, Fpg (*E. coli*) and OGG1 (human) (*13*).

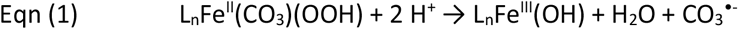

We therefore sought to identify the reactive oxygen species (ROS) formed in cells stressed with H_2_O_2_ when pre-equilibrated with 0-20 mM HCO_3_^-^. We assayed the metabolomes of *E. coli* and HEK293T cells, the RNA pool, and DNA lesions in HEK293T telomeres under these conditions to find that the metabolic product CO_2_ plays an important role in modulating and focusing the cellular Fenton reaction.

### Metabolic indicators show lower oxidative stress in cells as CO_2_/bicarbonate increases

HEK293T cells were initially grown under a standard 5% CO_2_ atmosphere (corresponding to 17 mM HCO_3_^-^) in DMEM medium to ∼80% confluency before their transfer to PBS with known concentrations of NaHCO_3_ (0, 5, 20 mM). Cells were given sufficient time to equilibrate between the medium and cytosol (1 h) following HCO_3_^-^ addition (*14*), and were then treated with 100 μM H_2_O_2_ for 15 min, a protocol that induced DNA damage at all four nucleotides indicative of HO^•^ when only 4 mM HCO_3_^-^ was present (*15*). Metabolomics profiling by HPLC-MS/MS conducted as a function of HCO_3_^-^ concentration showed that both the ratio of reduced glutathione (GSH) to its oxidized form GSSG (GSH:GSSG) and reduced NADH to NAD^+^ were smallest when oxidation reactions were conducted without added HCO_3_^-^, and greater levels of the reduced forms were found in cells with physiological levels of HCO_3_^-^ present (20 mM; Fig. 1A, 1B). These initial observations suggest that HCO_3_^-^ has a protective effect by minimizing the depletion of cellular antioxidants when H_2_O_2_ is the oxidant.

**Fig 1.**
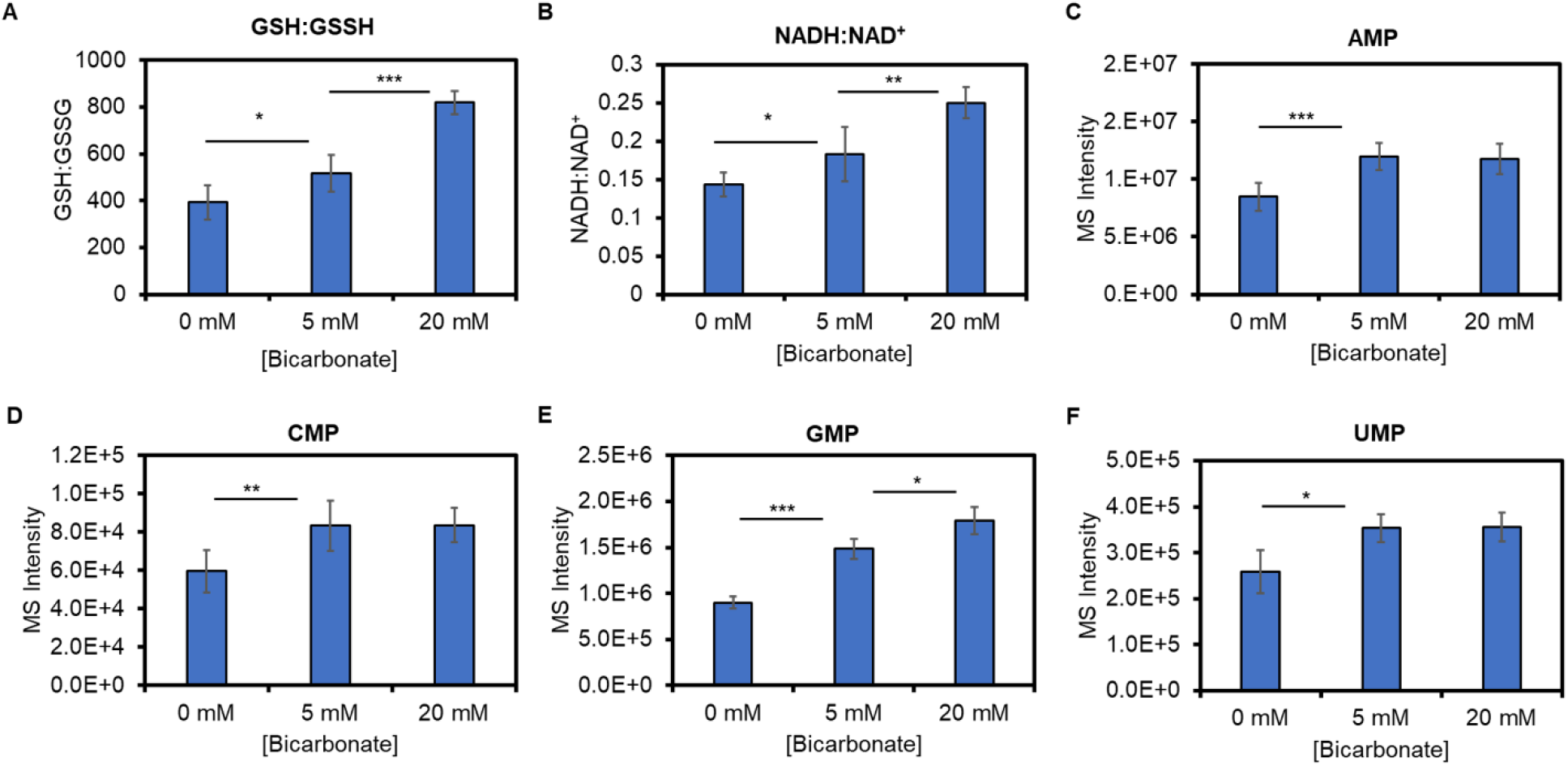
changes in redox-active metabolite concentrations after adding h_2_o_2_ to the medium with dependency on the bicarbonate concentration. hplc-ms/ms provided quantification of metabolite concentrations in hek293t cells pre-incubated in pbs medium with 0, 5, or 20 mm hco_3_^-^ for 1 h at 37 °c under atmospheric co_2_ before adding 100 μm h_2_o_2_. the reactions were allowed to progress for 15 min before quenching and analysis. ratios for reduced vs. oxidized states of (a) gsh:gssg and (b) nadh:nad^+^ are reported, and the mass spectrometry (ms) intensities are provided for the nucleotide-monophosphates (c) amp, (d) cmp, (e) gmp, and (f) ump. the analyses were conducted with 3-6 replicates and the levels of statistical significance are represented by * *p* < 0.05, ** *p* < 0.01, and *** *p* < 0.001 calculated by a student’s t-test.

Bicarbonate dependent changes to the nucleotide monophosphate (NMP) component of the metabolome provide clues regarding ROS identity. Without HCO_3_^-^ addition, all four NMPs were at their lowest levels after H_2_O_2_ oxidation (Figs. 1C-1F); all four NMP levels increased in the presence of 5 mM HCO_3_^-^, but only GMP, the nucleotide most sensitive to oxidation, further increased when 20 mM HCO_3_^-^ was present during oxidative stress (Fig. 1E). These data are consistent with bicarbonate changing the cellular Fenton reaction mechanism and generating a milder oxidant, CO_3_^•-^, rather than HO^•^, which is oxidizing only GMP.

### Apparent increase in 8-oxoguanosine levels in RNA is due to a focusing of oxidation on G

We profiled the transcriptomes of both *E. coli* and HEK293T cells for oxidation products and their formation sites. *E. coli* cells were grown in LB Miller medium under atmospheric CO_2_ (0.04%) to stationary phase, then incubated in PBS buffer with 0-20 mM HCO_3_^-^ and stressed with 100-μM H_2_O_2_ for 15 min, identically to the HEK293T cell studies. Total RNA was extracted in the presence of an iron chelator and an antioxidant to prevent artifactual oxidation (*16*).

Major products from RNA oxidation include 5-hydroxycytidine (ho^5^C), 5-hydroxyuridine (ho^5^U), 8-oxoadenosine (OA), and 8-oxoguanosine (OG), and these were measured before and after oxidative stress by HPLC coupled with UV and electrochemical (ECD) detectors (HPLC-UV-ECD). The nucleosides analyzed were liberated from total RNA by nuclease and phosphatase digestion. These four analytes allow profiling of known products of HO^•^ oxidation on each of the four nucleotides (*17*), are redox-active, and can be quantified with an ECD detector (*18*), which is a more sensitive method than MS/MS detection (*19*). This feature was needed because HCO_3_^-^ partially quenches the oxidation reaction (Fig. 1), leading to low overall levels of RNA lesions.

In *E. coli* exposed to H_2_O_2_ without HCO_3_^-^ present, significant increases in all four nucleoside biomarkers of HO^•^ oxidation were observed above background (Fig. 2A). With 5 mM HCO_3_^-^ present during the oxidation, ho^5^U and OG were formed significantly over background, and with 20 mM HCO_3_^-^, only OG was significantly formed above the background level while ho^5^C decreased below background (Fig. 2A). It should be noted that ho^5^C is enzymatically written into *E. coli rRNA*, contributing to its high background. Therefore, a decrease in ho^5^C upon oxidative stress may reflect its sensitivity to further oxidation or phenotypic changes akin to thermal and metabolic stress, as previously found for 23S rRNA ho^5^C2501 (*21*). The observation that G is the only nucleobase subject to oxidation by CO_3_^•-^ (*9*), supports a claim that physiological levels of HCO_3_^-^ impact the Fenton reaction to yield a milder one-electron oxidant, thereby oxidizing only the G heterocycle (Eqn. 1).

**Fig 2.**
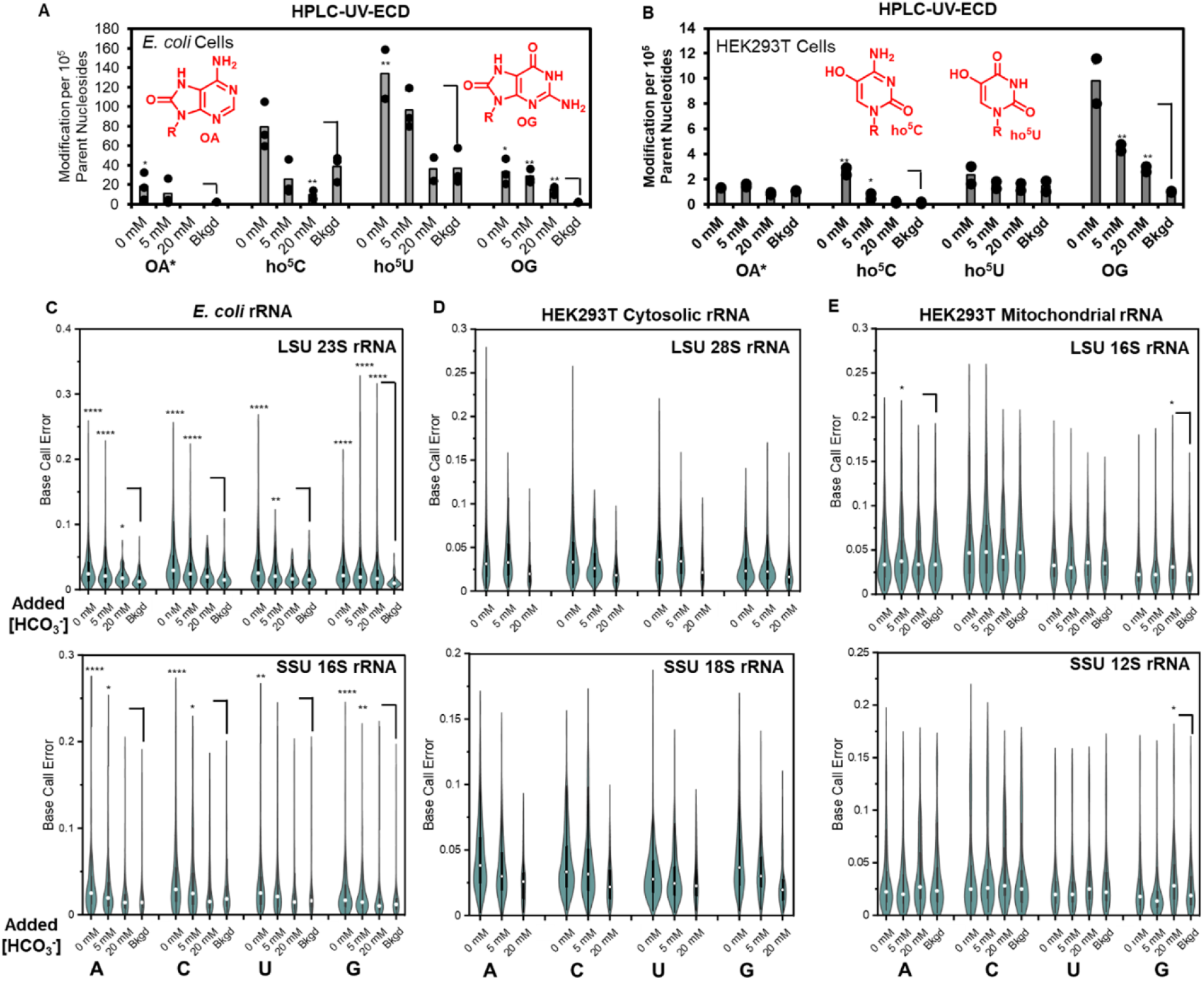
Profile of RNA oxidation products or sites upon adding H_2_O_2_ to *E. coli* or human cells showing dependency on the bicarbonate concentration. The redox-active products from RNA base oxidation ho^5^C, OA, ho^5^U, and OG were profiled in (A) *E. coli* or (B) HEK293T total RNA via nuclease/phosphatase digestion of the polymers to nucleosides and HPLC-UV-ECD quantification. RNA direct nanopore sequencing of the LSU and SSU rRNA from (C) *E. coli*, (D) HEK293T cytosolic rRNA, and (E) HEK293T mitochondrial rRNA were profiled from the oxidized cells to measure changes in base miscalls with ELIGOS2 that report on modification sites in the strands (*20*). The base miscall analysis for the *E. coli* and mitochondrial rRNAs was conducted by comparison of the cellular RNAs against a synthetic RNA of the same sequence without modification made by in vitro transcription (IVT). This approach permitted a profile of those RNAs in cells not exposed to oxidant (i.e., background (bkgd)). In contrast, the human cytosolic rRNAs are too G/C rich to allow synthesis of the RNAs without modifications via IVT; therefore, the comparison for the oxidized samples was against the rRNA from the non-oxidized cells resulting in no background being reported. Prior work and controls reported herein identified oxidation sites in RNA yield base call errors. The background for the *E. coli* was obtained from cells grown for 24 h at 37 °C in LB Miller medium under atmospheric CO_2_ levels, and the HEK295T cells were grown to ∼80% confluency in DMEM medium in a humidified incubator with 5% CO_2_ at 37 °C. The cells were placed in PBS with 0, 5, or 20 mM bicarbonate for 1 h (HEK293T) or 2 h (*E. coli*) before adding 100 μM H_2_O_2_. The oxidations proceeded for 15 min before quenching and harvesting the total RNA for analysis. The analyses were conducted in triplicate for HPLC-UV-ECD and duplicate trial for direct RNA nanopore sequencing with levels of statistical significance represented by * *P* < 0.05, ** *P* < 0.01, *** *P* < 0.001, and **** *P* < 0.0001 calculated by a student’s T-test. *The OA values from the HPLC-UV-ECD analysis are an overestimation resulting from A and OA coeluting in the HPLC.

In HEK293T cells, the RNA oxidation product levels were >3-fold reduced compared to *E. coli*, reflective of the evolved tolerance of human cells to oxidative stress compared to prokaryotic cells. Nonetheless, without HCO_3_^-^ present, ho^5^C and OG were significantly increased upon H_2_O_2_ addition, while OA and ho^5^U increased above the background but not significantly (Fig. 2B). The most striking feature of the oxidation products was that OG levels remained >2-fold above the background when 20 mM HCO_3_^-^ was present, while the other oxidative products were at background levels. These findings are consistent with the *E. coli* results and lead to the same conclusions: oxidation is focused on guanine when physiological bicarbonate is present.

Next, the extracted RNA strands before and after oxidation from both *E. coli* and human cells were analyzed by nanopore sequencing. This approach allows for global changes in RNA oxidation to be inspected both before and after oxidation without biasing the experiment by analyzing single oxidation products. A common approach to analyzing RNA direct nanopore data is to locate and quantify modifications in base-called data after reference alignment (*20, 21*). This approach to finding modifications works by searching for nucleotides with enhanced base miscalls, which arise because base calling was trained on canonical nucleotides, resulting in a higher potential of base miscalls at modified nucleotides (*20*). A drawback to this approach is the identity of the oxidation product is unknown; nonetheless, the nucleotide modified is known. As a consequence of rRNA comprising ∼80% of the total cellular RNA and its ease for high-depth sequencing, our analysis targeted the large and small subunit (LSU and SSU) rRNAs from the two cell types studied.

Nanopore sequencing of the SSU (16S) and LSU (23S) rRNAs from *E. coli* before and after H_2_O_2_-mediated oxidation in 0, 5, or 20 mM HCO_3_^-^ revealed that base-call errors increased upon oxidation with HCO_3_^-^ dependency, giving greater error frequencies (Fig. 2C). The 23S rRNA showed significantly greater base-call error at all four nucleotides when oxidations were conducted in 0 or 5 mM HCO_3_^-^. This observation is consistent with prior studies that found all four nucleobases could be oxidized at low bicarbonate concentration (*15*). With 20 mM HCO_3_^-^ present, oxidation led to the greatest increase in base-call error at G sites with a small increase at A sites, suggesting these nucleotides, particularly G, dominate the location of oxidation. Lastly, the base-call error for the SSU rRNA (16S) followed the same trend while being less intense compared to the results from the LSU, a finding consistent with a prior study inspecting H_2_O_2_-mediated oxidation of *E. coli* rRNA under only atmospheric levels of CO_2_ (*22*).

The cytosolic rRNAs in HEK293T cells when nanopore sequenced before and after the HCO_3_^-^-dependent oxidation followed a similar trend as found in *E. coli* (Fig. 2D). That is, without added HCO_3_^-^ all four nucleotides showed enhanced base-call error, and as the HCO_3_^-^ concentration was increased to 20 mM, the base-call error focused on G; moreover, the LSU rRNA (28S) was found to have greater error compared to the SSU rRNA (18S), similar to the *E. coli* results. The combination of product profiling and sequencing found oxidative stress imposed by H_2_O_2_ results in the least number of oxidatively derived lesions in RNA when physiological HCO_3_^-^ (20 mM) is present, and when damage does occur, it is focused on the G nucleotide.

### High bicarbonate levels in mitochondria buffer against oxidative damage

Eukaryotic cells utilize different ribosomes in the cytosol vs. mitochondria (*23, 24*); thus, we sequenced both types of rRNA before and after H_2_O_2_ oxidation. The mitochondrial rRNA generally displayed a non-significant change in base-call error before and after the oxidations throughout the added HCO_3_^-^ concentration series (Fig. 2E). A small increase in base-call error was found at G in both the SSU and LSU mitochondrial rRNAs when 20 mM HCO_3_^-^ was present during the bolus addition of H_2_O_2_. These data reveal that the compartmentalization of mitochondrial rRNA, and its different structure compared to the cytosolic rRNAs (*23, 24*), may be protective against oxidative stress.

### DNA damage in telomeres is focused on G when HCO_3_^-^ is present

To understand the nature and impact of Fenton chemistry on human genomic DNA damage, we focused on the telomere sequence 5’-(GGGTTA)_n_-3’/3’-(CCCAAT)_n_-5’ using a glycosylase-dependent, qPCR telomere length assay to quantify glycosylase-sensitive sites (Fig. 3A) (*25, 26*). Telomere length analysis before and after oxidation measures the strand breaks occurring during oxidation, a hallmark of HO^•^ chemistry (*17, 27*). The bifunctional glycosylase EndoIII yields a strand break at pyrimidine oxidation sites (e.g. thymine glycol, Tg) that can be quantified by the qPCR method (*28*), EndoIV yields strand breaks at abasic sites and oxidized abasic sites that are similarly quantifiable, and the bifunctional glycosylase Fpg creates strand breaks at purine oxidation sites, predominantly targeting G oxidation products such as OG (Fig. 3A) (*29*). The telomere sequence is balanced in A, C, G, and T content, allowing the relative reactivity of the nucleotides toward Fenton chemistry to be monitored at different levels of bicarbonate.

**Fig 3.**
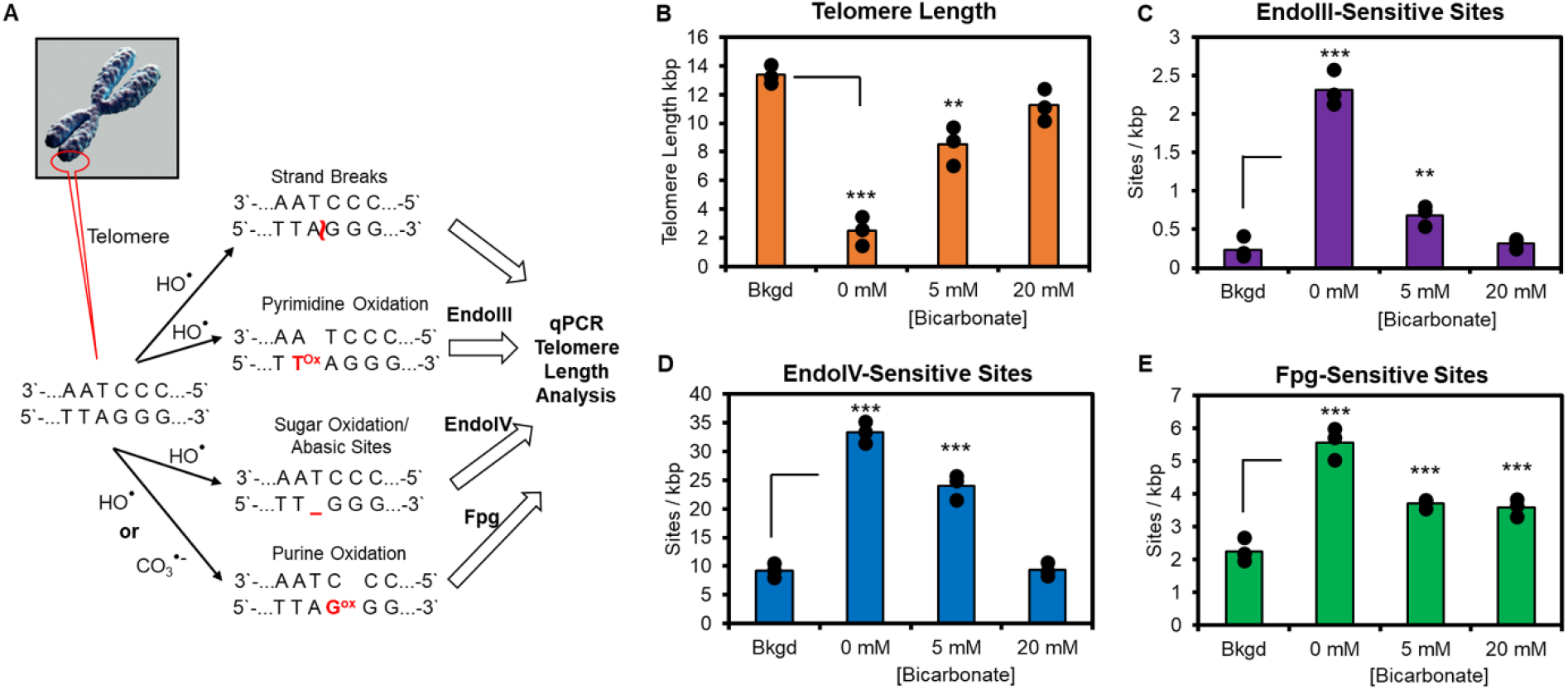
Bicarbonate dependency in telomere oxidation sites upon addition of H_2_O_2_ assayed by qPCR telomere length measurements. (A) Scheme to illustrate telomere DNA oxidation types analyzed via glycosylase removal of the damaged site. Direct measurement of the telomere length before and after oxidation reports on frank strand breaks that occur upon oxidation. Pyrimidine oxidation sites are revealed by EndoIII, an enzyme for which the preferred substrate is Tg, while the glycosylase can also remove 5hoC and 5hoU from DNA. Sites of sugar oxidation and abasic sites are substrates for EndoIV to yield strand breaks that are quantified by telomere-specific qPCR. Purine oxidation sites are found by Fpg, a glycosylase that favorably removes OG and Fapy-G from DNA, while the enzyme can also remove OA and Fapy-A. The bicarbonate dependency in H_2_O_2_-mediated oxidation of the telomeres was followed by qPCR to quantify (B) strand breaks, (C) EndoIII-sensitive sites, (D) EndoIV-sensitive sites, and (E) Fpg-sensitive sites. The oxidations were conducted by adding 100 μM H_2_O_2_ to HEK293T cells pre-equilibrated for 1 h in PBS medium with 0, 5, or 20 mM bicarbonate at 37 °C under atmospheric CO_2_ levels, followed by reaction quenching and harvesting of the gDNA. The background (bkgd) measurements were obtained from cells not exposed to oxidant. The analyses were conducted in triplicate with levels of statistical significance represented by ** *P* < 0.01 and *** *P* < 0.001 calculated by a student’s T-test.

We found that oxidative stress in the absence of HCO_3_^-^ led to the shortest telomeres, followed by a progressive increase in telomere length as the HCO_3_^-^ concentration increased (Fig. 3B). Inspection of HCO_3_^-^ dependency on EndoIII-sensitive sites after oxidation identified that without the protective effect of the buffer, substrates for EndoIII were maximal and decreased to background levels with physiological levels of HCO_3_^-^ (Fig. 3C). EndoIV-sensitive sites were also maximal without HCO_3_^-^ present and decreased to background with 20 mM HCO_3_^-^ equilibration before H_2_O_2_ addition (Fig. 3D). Finally, Fpg-sensitive sites, which predominantly report on G oxidation, were significantly increased at all three concentrations of HCO_3_^-^ studied (Fig. 3E). Collectively, the glycosylase-dependent telomere assays demonstrate significant damage in each case for oxidation when only 5 mM HCO_3_^-^ was present, consistent with a prior study using 4 mM HCO_3_^-^ and 100 μM H_2_O_2_ (*15*). A key difference in this study is that when the HCO_3_^-^ buffer is present at physiological concentrations, i.e. ≥20 mM, the DNA damage is focused on G nucleotides, as demonstrated by the series of glycosylase-dependent qPCR telomere length assays and the absence of frank strand breaks.

### DNA/RNA oxidation by H_2_O_2_/HCO_3_^-^ is iron dependent

We also explored whether iron functions as the catalyst in H_2_O_2_ degradation to yield ROS (HO^•^ and/or CO_3_^•-^) in cells. Briefly, HEK293T cells were incubated with the ferroptosis-inducing compound erastin to elevate labile pools of cellular iron (*30*). After a 16 h incubation in 2 μM erastin, the labile iron concentration was found to have increased 5.2x above that of native cells (Fig. 4A). The level of OG in the total RNA as measured by HPLC-UV-ECD showed a 4-fold increase after erastin treatment compared to cells with basal labile iron levels (Fig. 4B). Upon H_2_O_2_ oxidation with 0, 5, or 20 mM HCO_3_^-^ present, the OG levels were always greater in erastin-treated cells compared to those without treatment (Fig. 4B).

**Fig 4.**
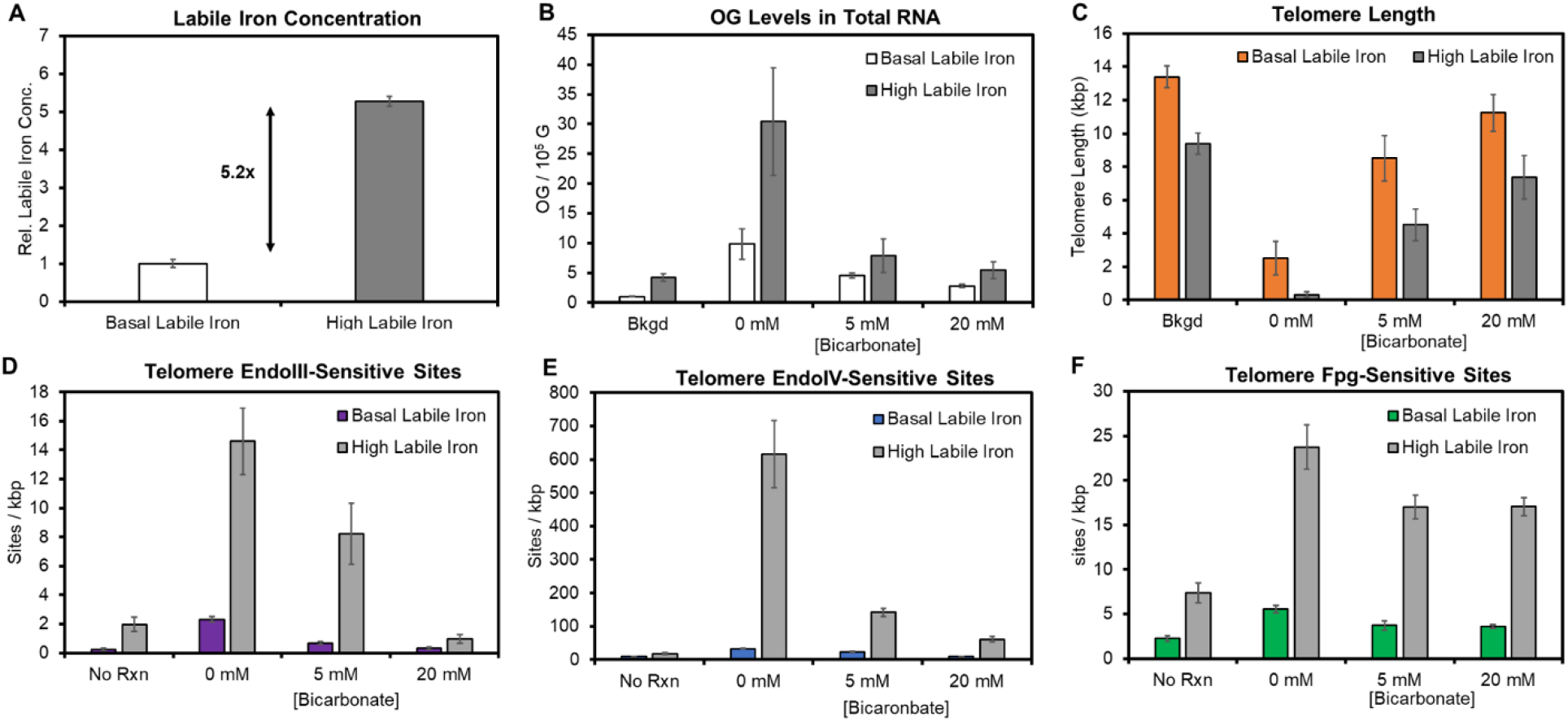
Iron dependency in oxidatively derived damage to HEK293T nucleic acids upon H_2_O_2_ addition. (A) Colorimetric assay determination of labile iron relative concentration before and after a 16 h treatment with the ferroptosis-inducing compound erastin. (B) Change in OG nucleoside levels in total RNA measured by HPLC-UV-ECD before and after erastin treatment, as a function of added H_2_O_2_ and bicarbonate to the culture medium. (C) Telomere lengths were measured by qPCR before and after erastin treatment and as a function of bicarbonate concentration during H_2_O_2_-mediated oxidation. qPCR quantification in telomeres of (D) EndoIII-, (E) EndoIV-, or (F) Fpg-sensitive sites in the high vs. basal iron level cells before and bicarbonate-dependent H_2_O_2_ oxidation. In all cases, background (bkgd) refers to cells grown under a 5% CO_2_ atmosphere in DMEM medium without exposure to H_2_O_2_. In the oxidation studies, the cells were equilibrated with 0, 5, or 20 mM bicarbonate in PBS for 1 h before adding 100 μM H_2_O_2_ for 15 min at 37 °C under atmospheric CO_2_ followed by quenching the reaction and harvesting the nucleic acids to be analyzed. The analyses were conducted in triplicate.

Telomeres in cells with elevated labile iron were assayed by the qPCR assay and compared to those with basal iron concentrations. When labile iron was elevated, telomere lengths were shorter than those found in cells with basal labile iron levels before and after each of the oxidation studies (Fig. 4C). The HCO_3_^-^ concentration dependency in telomere length measured after oxidation followed a similar trend in cells with basal or high labile iron levels (Fig. 4C). Moreover, EndoIII-sensitive sites in high labile iron cells were ∼6-fold greater than cells with normal labile iron levels under each condition studied (Fig. 4D). When 20 mM HCO_3_^-^ was present during the oxidation, even cells with high labile iron were protected from oxidation with the number of EndoIII-sensitive sites returning to the background level. The EndoIV-sensitive sites were also enhanced with high labile iron levels compared to basal levels (Fig. 4E); the enhancement in abasic and oxidized abasic sites was greatest for cells that lacked the protective effect of bicarbonate, whereas in the presence of 20 mM HCO_3_^-^, a ∼5-fold increase in EndoIV substrates remained, possibly reflecting the high level of DNA repair occurring in cells with elevated oxidative stress. In the final telomere analysis, Fpg-sensitive sites were consistently enhanced throughout the HCO_3_^-^ concentration series studied (Fig. 4F). When labile iron was increased, the telomere studies showed enhanced but nearly identical trends as those with basal iron, supporting Fe(II) as the catalyst for the Fenton reaction and highlighting the protective effect of physiological levels of HCO_3_^-^ buffer.

## Discussion

The small but reactive labile iron pool in cells is normally maintained at the Fe(II) state through removal of ROS by superoxide dismutase and catalase and by the presence of glutathione in low millimolar levels (*4*). Nevertheless, oxidative stress can be triggered by abrupt increases in metabolism, responses to inflammation, or the presence of redox-cycling natural products (*31*). Any resulting H_2_O_2_ is normally unreactive toward nucleic acids, but highly reactive with Fe(II), producing radical species capable of oxidizing DNA, RNA, and their nucleotide components (*32*). The literature to date argues that hydroxyl radical, or a related ferryl species (Fe=O^2+^), is the causative agent for scores of different oxidized nucleotide derivatives (Scheme 1) being formed from the cellular Fenton reaction. With such a broad range of reactivity, hydroxyl radicals would seem to inflict wide-ranging damage throughout the genome and transcriptome creating a nearly impossible task for DNA repair enzymes. In contrast, if the cellular Fenton reaction were focused on guanine oxidation, then a relatively small set of BER enzymes in *E. coli* and humans could act in response to oxidative stress, such as mutT/MTH1 to cleanse the nucleotide pool, fpg/OGG1 to remove OG:C base pairs in DNA, mutY/MUTYH for mutagenic OG:A base pairs, and the EndoVIII/NEIL glycosylases to remove hyperoxidized guanines from DNA and unhook oxidized crosslinks (*13, 33, 34*).

Indeed, a focus on guanine oxidation is precisely what we found for the cellular Fenton reaction. Metabolomics studies demonstrated that oxidative stress occurs when H_2_O_2_ is added to cell media, but the impact is reduced when HCO_3_^-^ is present. Here we find that a physiological relevant concentration of bicarbonate (20 mM) (*35*) is sufficient to redirect Fenton chemistry towards production of carbonate (CO_3_^•-^) rather than hydroxyl (HO^•^) radicals, as predicted by Meyerstein and coworkers (*6*). Importantly, CO_3_^•-^) is well documented to conduct one-electron oxidation reactions, especially for guanine, where the resulting G^•+^ can be hydrated and further oxidized leading to OG (*8, 9, 35*).

Beyond demonstrating the protective effects of CO_2_/HCO_3_^-^ and the focusing of oxidative modification on guanine, there are several noteworthy points from our DNA and RNA oxidation studies. For example, the nanopore sequencing studies directly compared human ribosomal RNA in the cytosol vs. mitochondria of the same cells. Cytosolic rRNAs are relatively G-C rich and include many solvent-exposed Gs on their surface (*36*). Accordingly, the addition of 20 mM HCO_3_^-^ before oxidative stress had a dramatic effect in lowering the base-calling error for all nucleotides (Fig. 2D). This same nanopore analysis for mitochondrial rRNA was somewhat different in that (a) the reactivity of G remains more obviously above background (Fig. 2E), and (b) there is less overall impact of adding 20 mM HCO_3_^-^ (Fig 2E). The latter point may indicate cells were still actively metabolizing and generating CO_2_ on their own in mitochondria. Further, estimates place the bicarbonate concentration in mitochondria near 100 mM (*37*)! This fact and our results may explain an earlier paradox in which the mitochondrial genome displayed less DNA damage than the nuclear genome (*38*), even though it is the organelle in which most H_2_O_2_ is generated.

A second key finding comes from the glycosylase-dependent qPCR studies of telomere oxidation. This assay, which can be performed on a small number of cells, clearly compares four different types of DNA damage: strand breaks, abasic sites, oxidized pyrimidines, and oxidized purines (Fig. 3A). Among these damage types, the first three return to baseline levels when 100 μM H_2_O_2_ is added as a stressor if 20 mM bicarbonate is present. Only guanine oxidation remains high in 20 mM HCO_3_^-^, showing that oxidative stress (15 min) is focused on G and is likely only marginally impacted by repair. Treatment with the ferroptosis-inducing compound erastin mimics these results, albeit at enhanced levels of DNA damage, supporting a conclusion that the oxidative damage induced by adding H_2_O_2_ is indeed iron dependent. Iron depletion via chelation before oxidative stress led to the expected inverse results.

The implications of these findings are broad, given the plethora of cellular pathways impacted by oxidative stress. Thousands of human genes are differentially expressed in response to a redox imbalance (*39*). Recent studies propose the oxidative modification 8-oxoguanine as an epigenetic base in promoters, where activating transcription factors such as NF-κB and HIF1-α can be recruited upon G oxidation (*40-42*). In such a mechanism, the cellular Fenton reaction focuses the oxidation on G where it impacts gene expression, and the modification is ultimately erased via the base excision repair pathway.

A confounding aspect of the oxidative DNA damage field is that ionizing radiation probably *does* generate hydroxyl radicals (*17*), leading to ribose and base chemistry as exemplified by the in vitro Fenton reaction lacking bicarbonate. In contrast, the present studies of the metabolome, the transcriptome, and the genome of both bacterial and human cells under H_2_O_2_ stress show unequivocally that the cellular Fenton reaction generates CO_3_^•-^, not HO^•^, when bicarbonate is present.

## ACKNOWLEDGMENTS

We acknowledge the Metabolomics Core Facility at the University of Utah Core (NIH-NCRR shared instrumentation grants 1S10OD016232-01, 1S10OD018210-01A1, and 1S10OD021505-01), and qPCR instrumentation from the HSC Genomics Core at the University of Utah. This work was supported by the National Institute of General Medical Sciences via grant no. R35 GM145237.

## Author contributions

Conceptualization and methodology: A.M.F. and C.J.B. Cell culture: A.M.F. and J.C.D. qPCR and nanopore sequencing: A.M.F. HPLC-UV-ECD and gel electrophoresis: J.C.D. Writing: A.M.F. and C.J.B. Supervision and funding acquisition: C.J.B.

